# Expansive Diversity and Temporal Dynamics of Emerging Polinton-like Viruses in a Marine Ecosystem

**DOI:** 10.64898/2026.05.28.728513

**Authors:** Ethan Mimick, Christopher Bellas, Nathan Ahlgren, Jed Fuhrman, Mohammad Moniruzzaman

## Abstract

Polinton-like viruses (PLVs) are an emerging group of double-stranded DNA viruses of microbial eukaryotes whose ecological roles in marine ecosystems remain poorly understood. Using a five-year monthly viromic time series from the San Pedro Ocean Time-series (SPOT), we investigated the diversity, temporal dynamics, and functional potential of PLVs in a coastal marine ecosystem. We identified 2,355 distinct PLV populations, revealing PLVs to be a highly abundant and diverse component of the marine virosphere. Phylogenetic analyses resolved multiple major PLV clades, including abundant Group X/Trimcap PLVs characterized by triplicate major capsid proteins, supporting the widespread occurrence of this unusual viral architecture in marine PLVs. Approximately half of PLV populations exhibited significant repeatable seasonal dynamics, partitioning into numerous chronotypes that reflect highly modular temporal niches. PLV abundance correlated positively with multiple productivity-linked environmental variables, including nitrate, particulate organic carbon, and primary productivity, suggesting close coupling between PLVs and seasonal ecosystem productivity. Functional analyses further identified diverse auxiliary metabolic genes in many PLV genomes involved in carbohydrate, lipid, redox, and nucleotide metabolism, with strong phylogenetic structuring across PLV clades. Together, these findings demonstrate that PLVs are abundant, functionally diverse, and ecologically dynamic members of marine viral communities, and suggest they are important yet underappreciated regulators of protist ecology and evolution in marine ecosystems.

## Introduction

Viruses are extraordinarily abundant in marine environments with ∼10 billion particles per liter of seawater with lifestyles as varied as the host they infect (Middelboe & Brussaard, 2017). Although substantial advances have been made on understanding the impact of viruses on marine prokaryotic communities, insights on their impacts on marine eukaryotes, specifically single celled eukaryotes (*aka* protists) remains fragmentary. Foundational ecological models such as *kill-the-winner* and *piggyback-the-winner* (Knowles et al., 2016; Winter et al., 2010) have provided frameworks for interpreting virus–host dynamics, with implications for life strategies of bacterial viruses ranging from classic predator-prey interactions to more persistent, less lethal consequences (Castledine & Buckling, 2024). Yet, these hypotheses are only beginning to be tested in the context of microbial eukaryote-virus interactions.

Microbial eukaryotes (also known as protists) make up a large portion of planktonic biomass and inhabit every corner of the ocean. These unicellular organisms have lifestyles as varied as the niches they inhabit. Their broad distribution and abundance makes them particularly important for nutrient cycling and overall geochemical cycling (Caron et al., 2017; Keeling & Campo, 2017; Yeh & Fuhrman, 2022c). Resolving how viruses regulate the population dynamics of marine protists thus remains of paramount importance. A key knowledge gap towards addressing this goal is that there still remains a large gulf between our understanding of the diversity and distribution of the marine virus groups infecting protists and their interaction strategies with their hosts.

Among recently discovered viruses that infect microbial eukaryotes, polinton-like viruses (PLVs) have emerged to be consequential given their environmental distribution and evolutionary relationship with mobile genetic elements that show predominantly transposable element-like lifestyle but harbor viral structural proteins (Koonin et al., 2024). PLVs belong to the broader polinton-like supergroup within the phylum *Preplasmiviricota* and are thought to be evolutionarily linked to not only polintons, but also virophages -parasites of giant virus replication resources. A recent study found PLVs to be particularly abundant in freshwater environments (C. M. Bellas & Sommaruga, 2021), while another (Piedade et al., 2024) study has found diverse PLVs in Antarctic waters - suggesting the presence of untapped diversity of PLVs in broader marine ecosystems. Genomically, PLVs are relatively compact double-stranded DNA viral elements, typically on the order of ∼14–40 kbp, that encode virion structural proteins such as major and minor capsid proteins along with replication, packaging, and integration-associated genes

PLVs exhibit an unusual suite of life strategies - including canonical lytic infection of single celled eukaryotes (Pagarete et al., 2015), and in some cases satellite/virophage-like interactions during giant virus co-infection, and diverse PLVs have been found to be integrating into the host genomes (C. Bellas et al., 2023; Bouchard et al., 2025; Koonin & Krupovic, 2017; Roitman et al., 2023; Thomas et al., 2026). In some systems, endogenous virophage-like elements can be activated during giant virus infection and reduce giant virus production, raising the possibility that some PLV-related elements may influence protist population dynamics not only through direct infection, but also by modulating giant virus-host interactions. These varied strategies and host species are of particular interest to marine ecology, as the diversity of life strategies displayed by PLVs make them a compelling group of viruses given their multifaceted potential to impact protist ecology and evolutionary trajectories. Therefore, it is critical to understand prevalence and dynamics of PLVs within marine ecosystems to fully resolve their roles in modulating the population dynamics of key protist lineages in the ocean.

Motivated by the knowledge gap on the diversity and ecological dynamics of polinton-like viruses in the ocean, we used a high temporal resolution, multi-year dataset from the San Pedro Ocean Time Series (SPOT) to conduct a systematic assessment of PLV diversity and temporal dynamics in a marine ecosystem. Specifically, the metagenomic dataset we leveraged were generated from the virome fraction from monthly water samples and were coupled with additional data on geochemical and oceanographic variables. This SPOT dataset covers a period of ∼5 years where monthly samples were collected - allowing for a first of its kind assessment of PLV seasonal dynamics over multiple years. Advances in metagenomics and recent developments in understanding the diversity of PLVs now allow robust *in silico* identification of PLVs (Stephens et al., 2024; Yutin et al., 2015), enabling us to explore their temporal patterns, genomic features, and putative ecological roles.

Our analysis revealed a remarkable diversity of PLVs in the SPOT dataset, suggesting that PLVs are a key group of protist viruses in the marine ecosystems, along with other recognized protist viruses, such as diverse RNA virus lineages and giant viruses (phylum *Nucleocytoviricota)*. Our analysis revealed novel PLV lineages, a substantial portion of which exhibit triplicate major capsid proteins. Assessment of their seasonality found ∼50% of the PLVs exhibiting significant, repeatable seasonal variations in their abundance profiles. We also identified several auxiliary metabolic genes (AMGs) in numerous PLVs, distribution of which is likely shaped by the phylogenetic history of these viruses. Together, these results reveal remarkable diversity and temporal dynamics of PLVs in marine systems, and provide strong evidence that PLVs are a key player in modulating protist ecology in the global ocean.

## Results

### Polinton-like virus diversity

We investigated the monthly virome time series (spanning five years) dataset from San Pedro Ocean Time Series (SPOT) to assess the diversity and temporal dynamics of polinton-like viruses (PLVs). This data consisted of 111 individual contig libraries, which we clustered to define distinct viral populations (see Methods). Within 2,355 of these populations, we identified major capsid protein (MCP) genes specific to PLVs. For each population, we selected the largest representative contig for downstream phylogenetic analysis. The length of the contigs representing the PLV populations ranged from 5,004 basepairs (bp) to 31,535 bp with an average length of 12,294 bp.

We performed phylogenetic analysis of MCPs encoded on population-representative contigs to assess the evolutionary diversity of PLVs at the SPOT site (Figure 1A). The MCP phylogeny revealed that a large proportion of SPOT PLVs segregated into four major clades, hereafter referred to as PLV1 through PLV4 (Figure 1). These clades primarily included PLVs encoding a single identifiable MCP homolog and therefore represent the canonical PLV diversity recovered from the SPOT dataset. The relative representation of these clades varied, with PLV2 forming the largest clade, comprising approximately 700 populations, whereas PLV1, PLV3, and PLV4 were smaller, each containing approximately 60 to 150 populations (Figure 1B). Notably, one subclade within PLV2 contained PgVV (Figure 1A), a previously described virophage-like PLV identified in association with the *Phaeocystis globosa* giant virus PgV-16T(Roitman et al., 2023). In addition to these major PLV clades, we recovered a small number of virophage populations, with only 28 populations assigned to virophages (Figure 1B), indicating that the SPOT dataset was dominated by PLV rather than virophage diversity.

**Figure 1.**
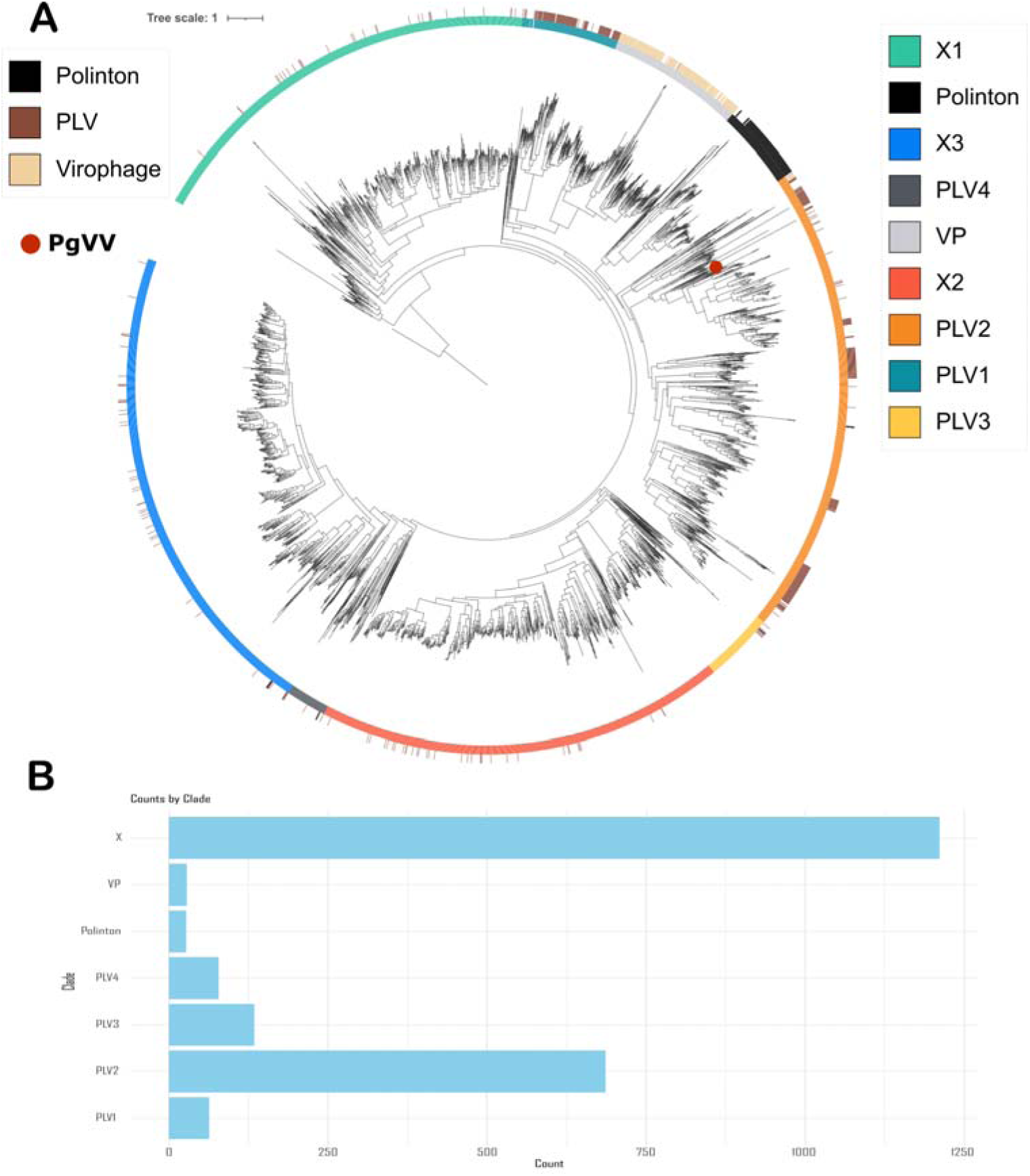
Phylogenetic classification of SPOT PLV populations. (A) Maximum likelihood major capsid protein (MCP) phylogeny showing SPOT PLV populations in relation to reference PLVs, polintons, and virophages. Colors on the outer ring indicate reference virus sequences, while the inner ring indicate major PLV clades identified in this study, along with a small subset of sequences belonging to the Polinton and virophage groups. (B) Bar plot showing the number of SPOT PLV populations assigned to each clade, highlighting the dominance of X group and PLV2 lineages in this dataset.

Beyond these canonical PLV clades, the MCP phylogeny revealed an unusual and highly represented group of PLVs that frequently encoded multiple MCP homologs, with many contigs harboring three distinct MCPs (Figure 1A). Importantly, MCPs from these multi-MCP PLVs did not scatter broadly across the PLV phylogeny. Instead, they resolved into three discrete phylogenetic clusters (X1, X2 and X3), and the three MCP copies encoded within individual PLV contigs did not group as sister clades (Figure 1A). This pattern argues against recent within-genome MCP duplication and instead supports the presence of three evolutionarily distinct capsid lineages that have been stably maintained within this PLV group. Approximately 1,200 PLV contigs encoded multiple MCPs and exhibited this phylogenetic pattern. Yutin et al 2015 first reported two PLVs encoding three distinct MCP homologs in metagenomic data, similar genomes were later found as a minor component of alpine freshwater lake viromes and termed Tri-major capsid or Trimcap PLVs (Bellas et al., 2021). More recently, Piedade et al. described an Antarctic PLV group, referred to as group X, in which multiple MCPs appeared to be a hallmark feature.

In contrast to these earlier reports, this Trimcap/group X-like lineage accounted for more than half of the PLV populations recovered from SPOT, suggesting that Trimcap-like PLVs are not rare outliers in this marine system but instead represent a prominent and evolutionarily distinctive component of oceanic PLV diversity.

### Environmental factors shaping PLV dynamics

A number of environmental variables positively correlated with total PLV relative abundance (RPKM), which included total bacteria, nitrite, nitrate, photosynthetically available radiation (PAR) saturation, particulate organic carbon, and primary productivity (Mantle correlation p-value<=0.05) (Figure 2). We found PLV1 clade to be positively correlated with nitrite, phosphate, PAR saturation, salinity, and temperature. PLV2 was positively correlated with absorbance due to phytoplankton, chlorophyll *a* saturation, chlorophyll maximum depth, phosphate, particulate organic carbon, primary productivity, salinity, and temperature (Figure 2). PLV3 and virophage populations were not correlated with any of the abiotic factors recorded, while PLV4 was positively correlated with chlorophyll a saturation, chlorophyll max depth, and salinity. Trimcap/group X PLVs were positively correlated only with oxygen. Many of the significant correlations involved variables linked to nutrient availability, light conditions, and water column structure (Figure 2). However, the strength of associations varied across PLV clades, with some lineages showing multiple environmental correlations while others showed little or none. These patterns indicate that PLV clades differ in their environmental responsiveness, consistent with ecological differentiation across evolutionary lineages.

**Figure 2:**
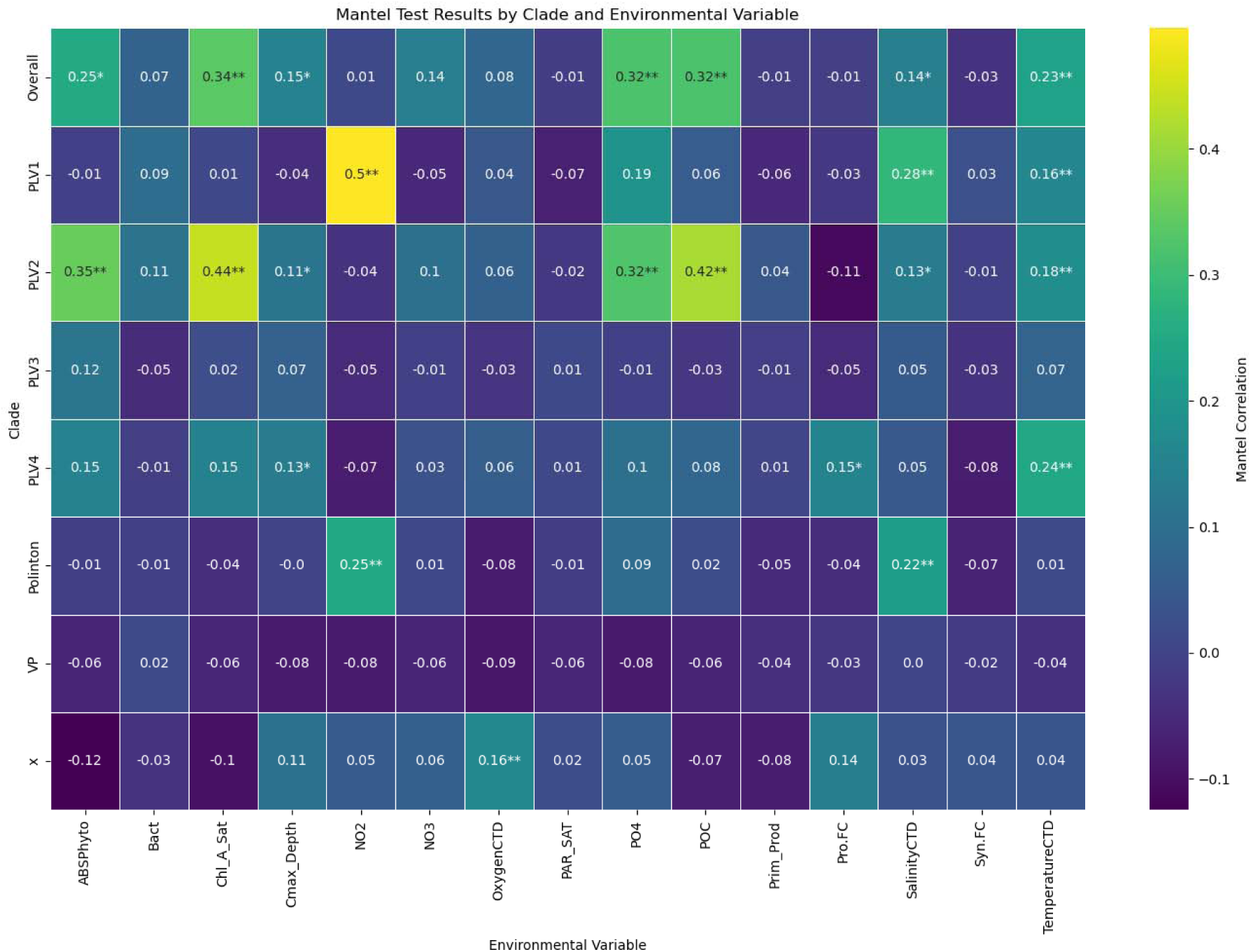
Environmental associations of PLV clades. Heatmap of Mantle scores for correlations between PLV clades and environmental variables. Rows show PLV clades, columns show environmental variables, and cell values indicate Mantel correlation coefficients. Color intensity reflects correlation strength and direction, and asterisks denote statistical significance (*: p<0.05, **: p< 0.01). Distinct correlation patterns among clades suggest that PLV lineages are associated with different environmental gradients. ABSPhyto : Absorbance due to phytoplankton, Bact : bacterial abundance, Chl_A_Sat: chlorophyll A saturation, Cmax_Depth : Chlorophyll maximum depth, NO2 : Nitrite, NO3 : Nitrate, OxygenCTD : oxygen level measured by CTD, PAR_SAT : photosynthetically available radiation saturation, PO4 : phosphate, POC : particulate organic carbon, Prim_Prod : primary productivity, Pro.FC : total Prochlorococcus measured by flow cytometry, SalinityCTD : salinity measured by CTD, Syn.FC : total Synechococcus measured by flow cytometry, TemperatureCTD : temperature measured by CTD.

### Seasonal Trends in Viral Abundance

We assessed seasonal trends in PLV population dynamics at SPOT based on the presence of significant variance in seasonal abundance of a population as compared to the mean abundance of all populations. This analysis revealed two distinct clusters of PLVs (termed cluster 1 and 2) - cluster 1 consists of the populations with clear seasonal signals, and cluster 2 represents the PLVs with dynamic abundance over time with no clear seasonal signals (Figure 3A, Supplementary figure 1).

**Figure 3:**
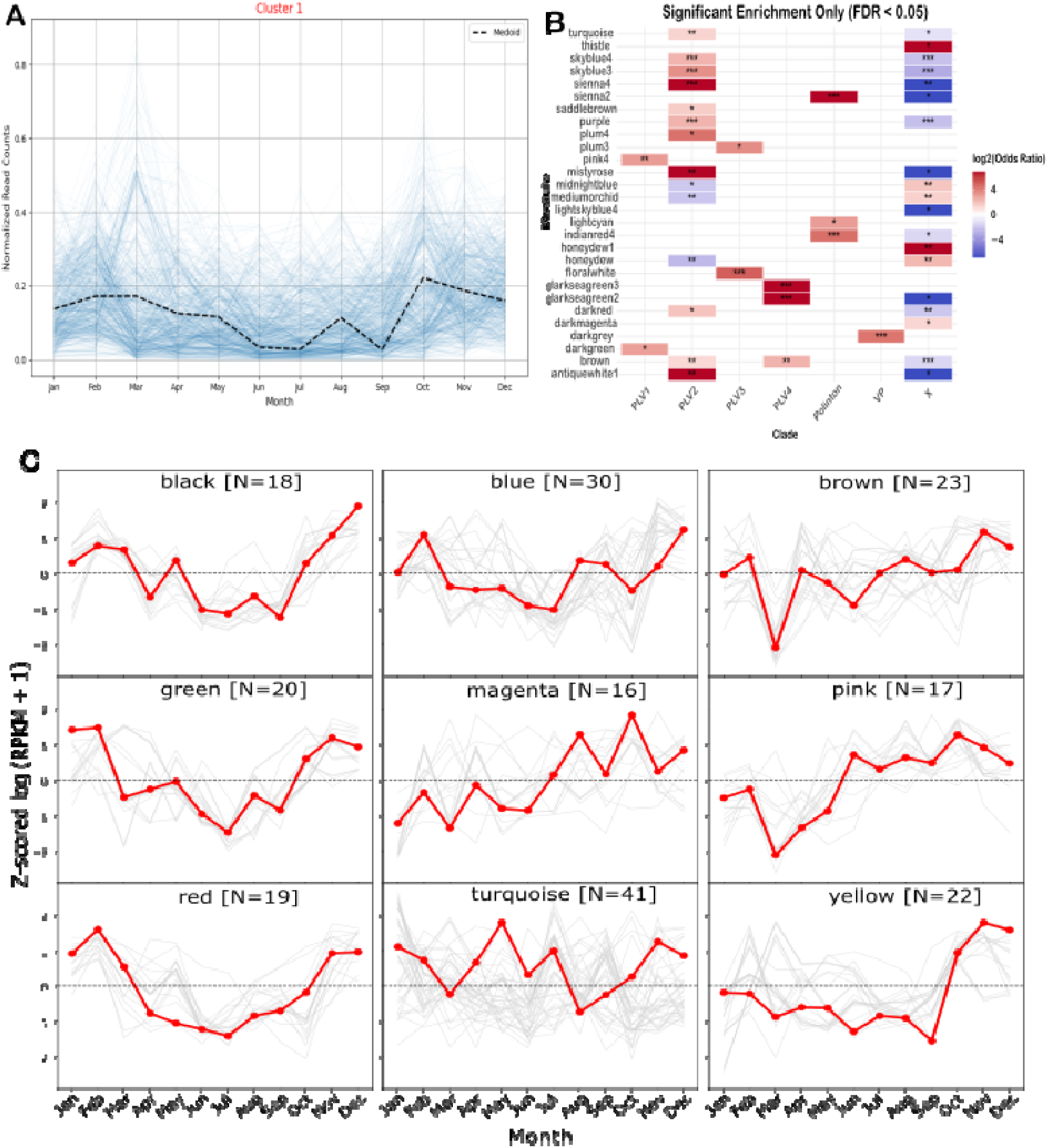
Seasonal patterns of SPOT PLV populations and their clade-level structure. (A) Median seasonal profile of a major seasonal cluster, summarizing the dominant temporal signal across populations. (B) Clade enrichment heatmap showing which PLV lineages are significantly over- or under-represented within seasonal modules (*: p<0.05, **: p< 0.01, ***: p<0.001). Together, these analyses show that SPOT PLVs exhibit diverse seasonal chronotypes and that temporal dynamics are partly structured by PLV phylogenetic identity. (C) Monthly abundance profiles of significantly seasonal PLV populations grouped into WGCNA modules. Gray lines represent individual PLV populations, while red lines show the average seasonal trajectory for each module, revealing distinct annual abundance patterns among PLV groups.

The seasonal cluster (cluster 1) contained 621 representatives. Among the seasonal PLV genomes identified at SPOT, WGCNA co-occurrence network resolved a large number of co-varying modules (chronotypes), indicating that seasonality is not driven by a single dominant temporal pattern but instead by many distinct, recurrent temporal trajectories distributed across the year (Figure 3B, C). The largest assigned modules included turquoise (41 genomes), blue (30), brown (23), yellow (22), green (20), red (19), black (18), pink (17), and magenta (16) (Supplementary table 1), revealing a highly structured but heterogeneous seasonal assemblage (Figure 3B). Together, these results suggest that seasonal PLV dynamics at SPOT are both recurrent and highly modular, with different subsets of genomes tracking distinct annual niches.

Inspection of the largest modules shows that several recurring seasonal strategies are present (Figure 3C). Some modules are characterized by late autumn to winter maxima. For example, the yellow module rises sharply in October through December, suggesting a tightly constrained late-year seasonal window. The black module shows a clear recovery in autumn and reaches its highest values in November and December. The brown module follows a related pattern, with strongest positive values in late fall and early winter, although with a pronounced dip in March. Other modules display broader cool-season enrichment. The green and red modules are elevated in January and February, decline strongly through spring and summer, and then recover again in late fall, indicating bimodal cool-season preference. Together, these patterns suggest that a large fraction of the more abundant PLV modules are centered on late-fall to winter dynamics. However, this seasonal structure is not exclusively winter-associated. The magenta and pink modules are more characteristic of warm-season to early-fall dynamics, with positive increases emerging from summer onward and strongest values in late summer or autumn. The blue module shows an intermediate pattern, with reduced abundance in spring to early summer and higher values in late summer and winter. Smaller modules further support this broader seasonal heterogeneity. Although many of the larger modules peak during winter or the late-year transition, several smaller modules show maxima outside this window, including spring-associated modules and a smaller but meaningful set of summer-associated modules, such as grey60 and purple (Supplementary Figure 2-4). Thus, PLV seasonality appears to be strongly biased towards cool-season and late-year abundance peaks, but not limited to them.

The module-by-clade enrichment analysis suggests that phylogeny contributes to these seasonal patterns, but only partially (Figure 3B). We identified 28 chronotypes disproportionately representing specific PLV clades, suggesting that the seasonal patterns of certain PLV clades were likely influenced by their phylogenetic history. In particular, multiple modules are enriched for PLV2, including several of the larger modules, while PLV4, Polinton, and virophages (VP) show enrichment in a smaller number of more specific modules (Figure 3B). This supports the idea that at least some components of seasonal recurrence are conserved within evolutionary lineages. However, the enrichment is often distributed across many modules, and the same broad clade can be associated with multiple distinct temporal patterns. In addition, we also found significant depletion of some clades from particular modules, reinforcing that lineage membership does not fully determine seasonal dynamics (Figure 3B). Because each module contains only a subset of the total seasonal PLV pool, these enrichments should be interpreted cautiously, but overall, they are consistent with a model in which phylogenetic relatedness partially constrains seasonal ecological dynamics.

### Functional Diversity of Polinton like Viruses

The PLVs commonly encoded the canonical replication and structural genes including ATPases, Major capsid (MCP) (including group X genomes with triplicate MCP genes), and to a lesser extent minor capsid (mcp) genes (Figure 4A, 5A). Apart from the genes involved in core replicative and structural functions, we also identified genes with possible metabolic functions in a substantial proportion of genomes. Specifically, 289 genomes encoded carbohydrate transferase domains, including galactosyl- and glycosyltransferases, suggesting that many PLVs possess the potential to manipulate host carbohydrate metabolism or glycan structures during infection (Figure 5B). These domains were more common in Trimcap (258 genomes), with smaller numbers present in PLV2 (29 genomes) and PLV3 (2 genomes). We also detected lipid metabolism–related genes in 13 genomes, which contain lipase (class 3) domains, indicating that a subset of PLVs may influence host lipid composition or membrane dynamics during infection. In addition, ribonucleotide reductase (RNR)- a metabolic enzyme commonly encoded by bacteriophages and giant viruses to support nucleotide biosynthesis during replication (Harrison et al., 2026; Sakowski et al., 2014) - was identified in 51 PLV genomes. Notably, these genes were more common in the PLV2 clade (37 genomes), with smaller numbers present in PLV1 (9 genomes) and only a few occurrences in VP (3 genomes) and group X (2 genomes), indicating a lineage-specific bias in nucleotide metabolism (Figure 5B). Other auxiliary metabolic genes occurred at lower frequencies, including nucleotide-diphospho-sugar transferases (9 genomes), mannosyltransferases (3 genomes), sulfotransferases (3 genomes), and UDP-galactopyranose mutases (2 genomes), along with glutaredoxin (2 genomes) and a single phospholipase A2–like enzyme. These genes suggest additional capacities for modifying host carbohydrate, lipid, and redox metabolism. We also assessed clade-specific functional annotations, defined as Pfam domains found exclusively within a single PLV clade (Figure 5A). Examination of presence–absence patterns of Pfam-annotated functions across PLV groups revealed that a large proportion of domains were restricted to a single clade. PLV2 contained the highest number of unique Pfam domains (150), followed by Trimcap (143) and VP (130) (Figure 5A). In contrast, only 25 functional domains were shared across all groups, and the greatest functional overlap occurred between VP and PLV2, which shared 64 domains. t-SNE clustering of genomes based on their functional profiles revealed clear grouping of PLV genomes according to their phylogenetic relationships (Supplementary Figure 5). In particular, genomes belonging to clade X and PLV2, the two largest clades, form distinct clusters, indicating that different PLV lineages possess distinct gene repertoires (Supplementary Figure 5). Together, these patterns suggest lineage-specific functional differentiation shaped by the evolutionary history of PLVs.

**Figure 4:**
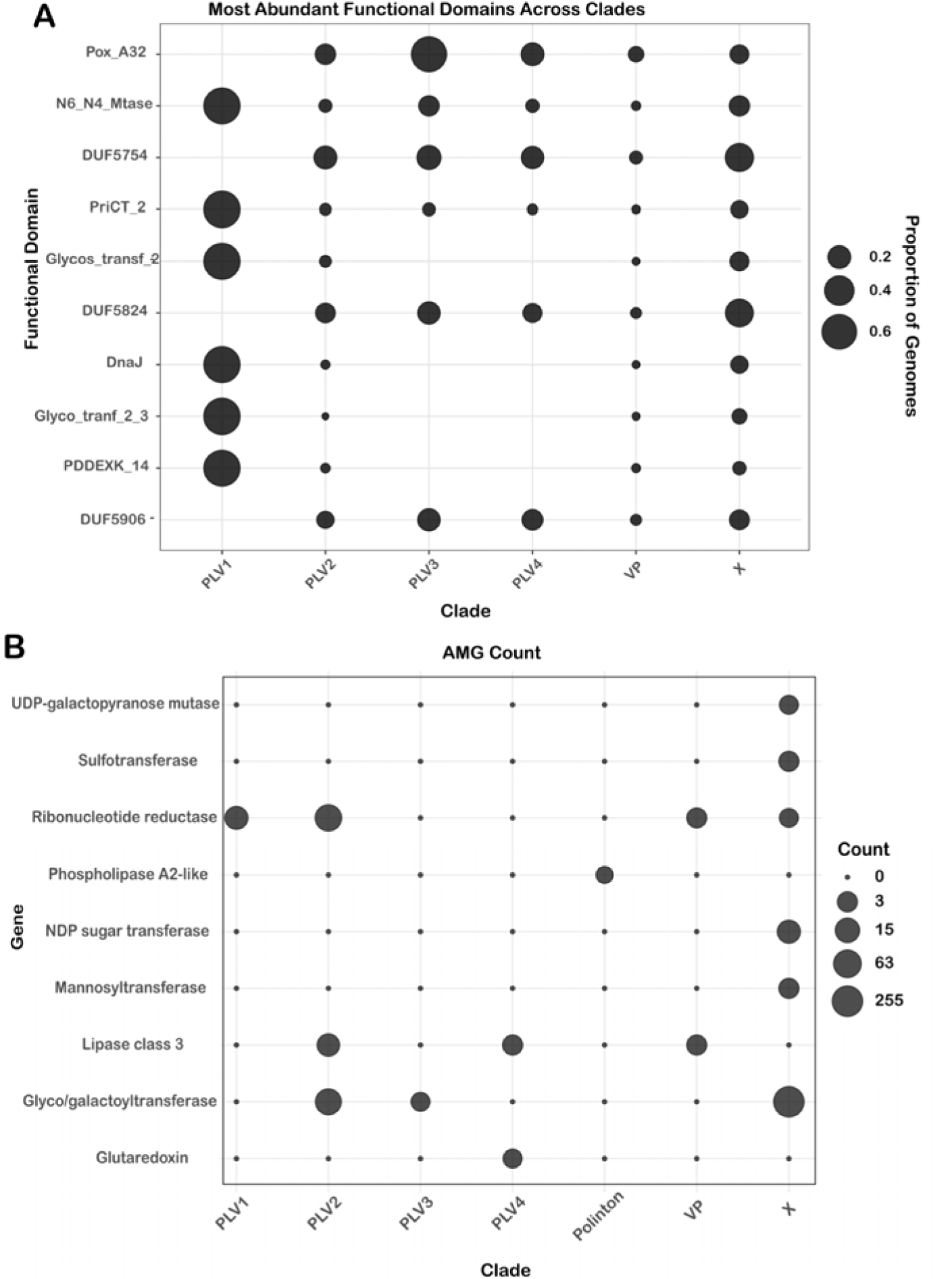
Functional profiles of SPOT PLV populations. (A) Bubble plot showing the ten most abundant functional domains across PLV clades, normalized by the number of genomes in each clade. Bubble size represents the proportion of genomes containing each domain. Pox_A32: Poxvirus A32-like ATPase, N6_N4_Mtase: N6/N4 Methyltransferase, DUF: Domain of unknown function, PriCT_2: Primase C-terminal, Glycos_transf: Glycosyl transferase, DnaJ: DnaJ/HSP40 family protein, PDDEXK_14: PD-(D/E)XK phosphodiesterase. (B) Bubble plot showing the distribution of putative auxiliary metabolic genes (AMGs) across PLV clades, with bubble size representing gene counts (log2 converted values). Together, these panels show that SPOT PLV clades differ in both core functional domain composition and metabolic gene content, suggesting lineage-specific variation in functional potential.

**Figure 5:**
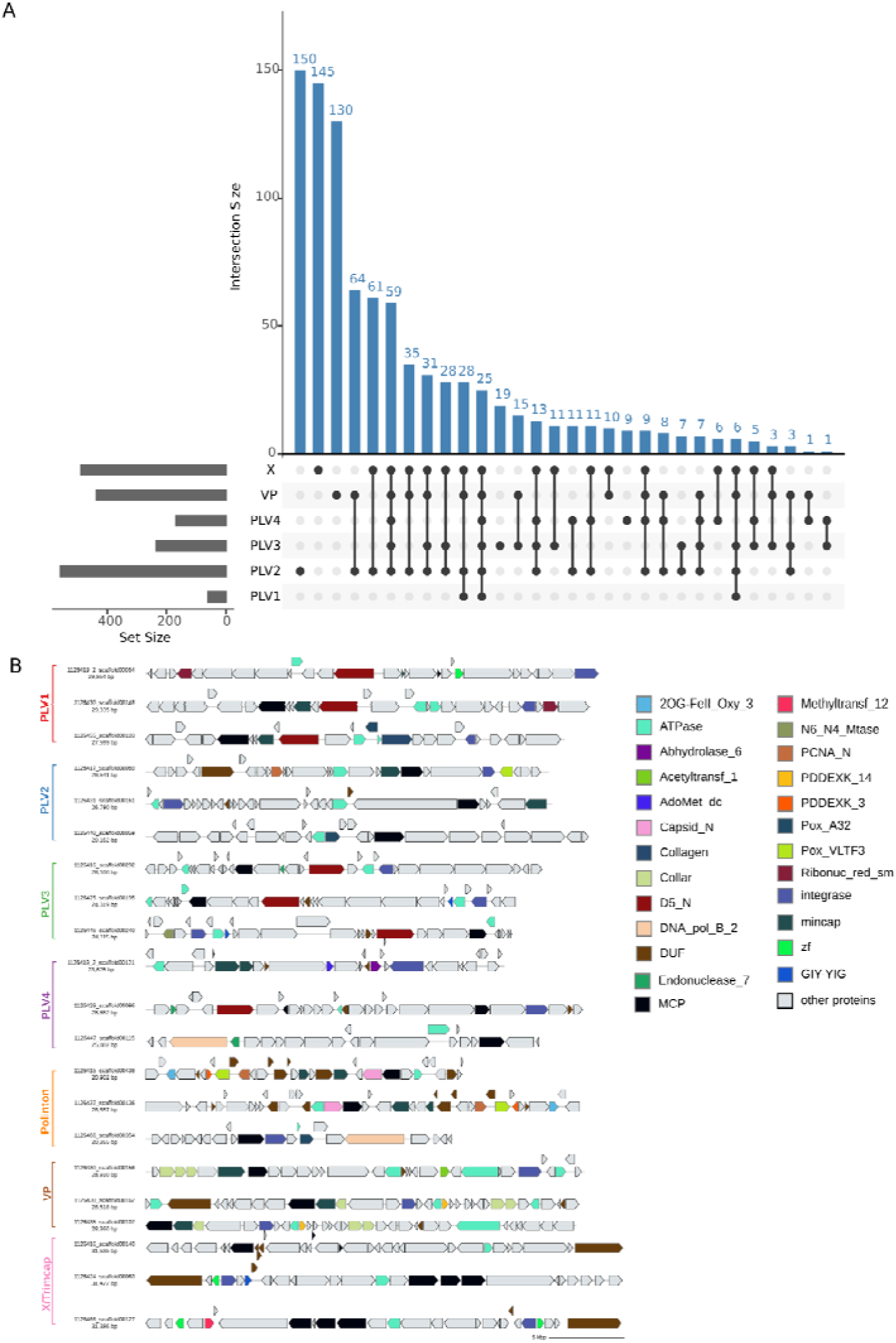
Patterns of shared and unique functions and genome architecture of SPOT PLV populations. (A) UpSet plot showing shared and unique functional domains across major SPOT PLV clades. Vertical bars indicate the number of domains found in each clade or clade combination, while horizontal bars show the total number of detected domains per clade. (B) Representative genome maps for each PLV clade, showing the organization of predicted genes, hallmark viral genes, and annotated functional domains. This comparison highlights a shared functional backbone among SPOT PLVs, while also revealing substantial clade-specific variation in gene content and genome architecture.

### Association of SPOT PLVs with co-occurring eukaryotes

Correlation of PLV abundance profiles with eukaryotic abundance profiles derived from 18S rRNA data yielded a co-occurrence network (Pearson’s r > 0.4), putatively linking PLVs to a diverse set of eukaryotic taxa at this site, spanning 12 distinct eukaryotic groups (Figure 6). The correlation strengths between eukaryotic and virus groups range from 0.91 to 0.4. Some notably strong connections are between a PLV2 and an ochrophyta (Pearson’s *r* = 0.915), group X and a dinoflagellata (*r =* 0.745), and PLV4 member with a ciliophora (*r=* 0.733) (Supplementary dataset 1). Overall, the network indicates that multiple PLV lineages are associated with a broad range of putative hosts. Trimcap/X group PLVs show particularly wide connectivity, including associations with dinoflagellates, ochrophytes, and ciliophora. Notably, they also exhibit repeated connections to metazoa, with three group X viruses connecting to the same metazoan node with moderate to strong correlation values (*r=*0.77, 0.71 and 0.56) (Supplementary dataset 1). PLV2 displays associations with dinoflagellates, ciliophora, and radiolaria, while PLV4 is also linked to metazoa and ciliophora. In contrast, more restricted patterns are observed for other groups, with PLV1 forming two connections to ciliophora and one to chlorophyta, and two PLV3 members showing connections to ochrophyta. One of the SPOT PLVs closely related to polintons was found to be associated with choanoflagellates, close relatives of animals (Figure 6).

**Figure 6:**
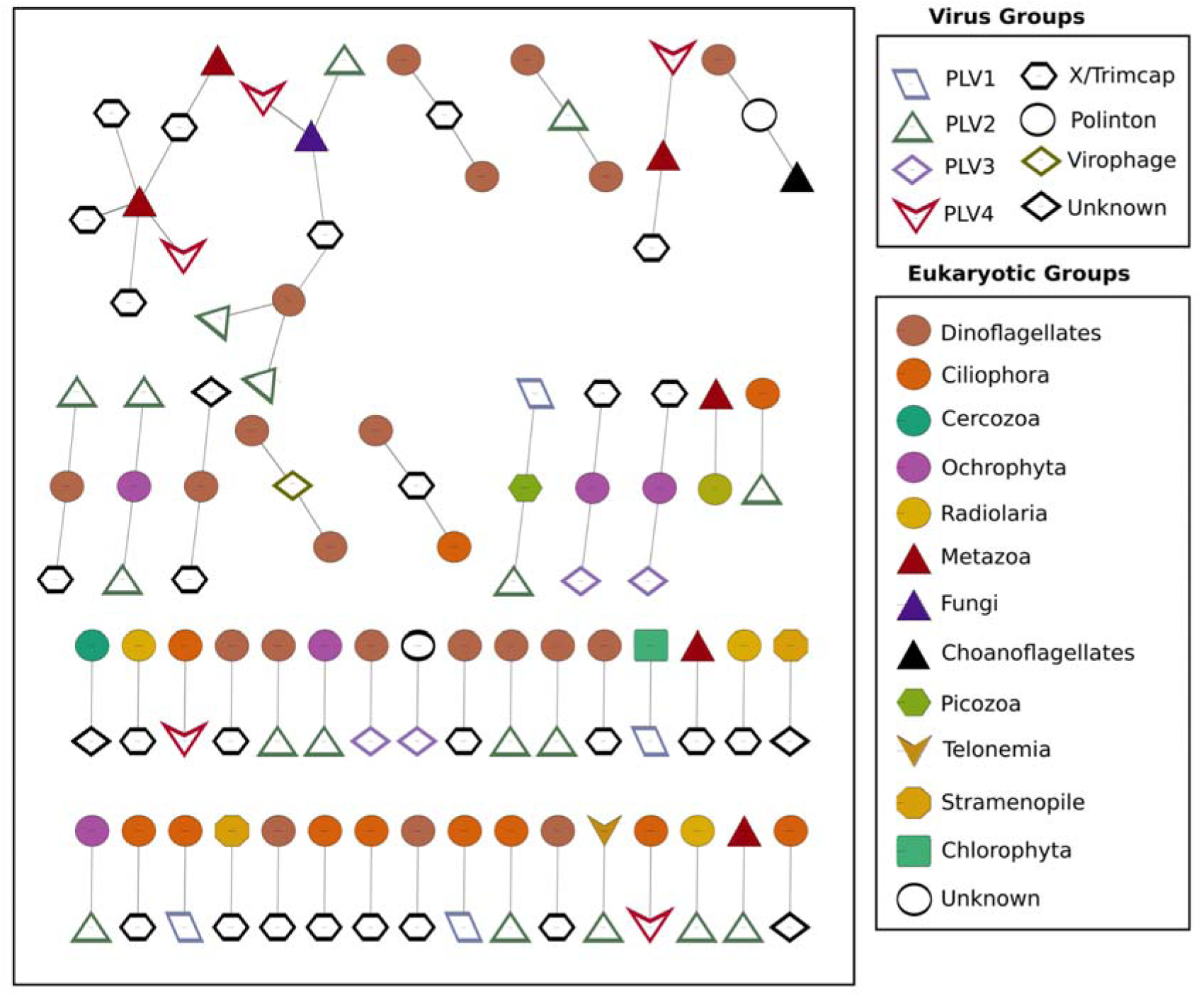
Correlation network of PLV members and co-occurring eukaryotic taxa in the SPOT time series. The network was built using FlashWeave and only connections with Pearson’s correlation >0.4 were retained. The network shows putative (but not definitive) host associations of certain PLV members. Also see supplementary dataset 1.

## Discussion

### Marine PLV diversity at SPOT

This study provides a multi-year ecological view of polinton-like viruses (PLVs) in a coastal marine ecosystem and shows that PLVs are not rare or sporadic components of the virosphere, but instead represent a highly diverse, and seasonally structured assemblage of eukaryotic DNA viruses. Phylogenetic analysis revealed a number of clades of PLVs in the SPOT site, with PLV2 and X group PLVs as the most numerous, consisting of 84% of the genomes we identified in this site. This seems to parallel an emerging pattern for giant viruses (large DNA viruses) of protists, which are dominated across the global oceans by two groups, Algavirales and Imitervirales, with other orders having reduced representation (Endo et al., 2020; Minch & Moniruzzaman, 2025; Moniruzzaman et al., 2020). Interestingly, the only other marine study on PLVs from a recent Antarctic study found Trimcap/X PLVs with unusual tri-capsid architecture to be the most abundant group of PLVs, which is consistent with what we found in our study (Piedade et al., 2024). This suggests that group X PLVs might be a key group of PLVs in the marine ecosystem. While the other groups had disproportionately fewer members, further studies are needed to better assess the distribution of diverse PLV clades in different oceanic regimes.

One of the key outcomes of this study is the confirmation and broader characterization of Trimcap PLVs carrying three phylogenetically distinct major capsid protein (MCP) homologs within a single genome. Although tricapsid PLVs were noted previously in freshwater systems by Bellas et al. and similar signatures were later reported from the Antarctic study, our phylogenetic analysis offers a clearer view of the origin of this architecture (C. M. Bellas & Sommaruga, 2021; Piedade et al., 2024). The three MCPs encoded by individual Group X genomes do not form monophyletic clusters. Instead, each of the three MCP homologs creates distinct phylogenetic groups in contrast to genome specific clusters which would be expected in case of recent genome-specific duplicates. Although direct functional evidence is still lacking, retaining these tri-capsid structures might contribute to complexity in the virion architecture or expanded host recognition capacity. The recurrence of the tri-capsid PLVs across geographically (Antarctic vs SPOT) and environmentally distinct datasets (marine and freshwater) suggests that multi-MCP PLVs have a broad ecological distribution.

The finding that virophage-like populations (Maveriviricetes) are relatively rare at SPOT(comprising only 28 populations) is particularly informative. Virophages are obligate parasites of giant virus replication compartments (La Scola et al., 2008; Nino Barreat & Katzourakis, 2023) and their scarcity here is intriguing given the fact that giant viruses are diverse and abundant in the SPOT sampling site. The evolutionary proximity of PLVs to virophages, established through shared protein fold structures and phylogenomic analysis (Koonin et al., 2015), raises the possibility that some PLVs at SPOT may occupy related ecological roles during giant virus infection. Indeed, previous research revealed that some PLVs can parasitize cellular or viral resources during giant virus replication (Roitman et al., 2023; Stough et al., 2019). In this context, the placement of PgVV within a PLV2 subclade is especially notable. PgVV is associated with the *Phaeocystis globosa* giant virus system (Roitman et al., 2023), linking the dominant PLV2 clade at SPOT to a lineage with known giant virus-associated biology. Although phylogenetic placement alone does not establish functional interaction, the presence of PgVV within PLV2 links this dominant SPOT clade to a lineage with known giant virus-associated biology and suggests that PLV2 may include ecologically relevant PLVs involved in algal host–giant virus systems.

### Ecological dynamics of PLVs at SPOT

Our study revealed a highly dynamic population of marine PLVs that responds to diverse environmental factors, supporting a view that a substantial fraction of SPOT PLVs are linked, directly or indirectly, to seasonal primary production. Several positive correlations involved variables associated with nutrient supply, and water-column productivity, including nitrate, nitrite, particulate organic carbon, PAR, chlorophyll-related variables, and primary productivity. Accordingly, the positive relationship between PLV abundance and production-linked variables is consistent with the possibility that many PLVs are associated with actively growing or seasonally recurring protist hosts. This interpretation is also supported more broadly by work on marine giant viruses and phages in the SPOT site done previously (Dart et al., 2023; Laperriere et al., 2025), which has linked viral dynamics to cyanobacteria or photosynthetic microbial eukaryotes that occupy seasonally shifting ecological niches. This is also consistent with the broader, well-documented seasonality of the whole microbial community at SPOT (Cram et al., 2015; Yeh & Fuhrman, 2022a). A curious departure in those patterns, however, is the frequent number of PLV populations that are more abundant in late fall/winter as opposed to spring/summer, whereas, for example, cyanophage populations were more evenly divided between these two broad chronotype patterns, generally connected to distinct host taxa (Dart et al., 2023). This could perhaps point to a major difference in the general ecology of PLVs that the seasonally driven populations are more frequently associated with processes with colder, winter months and many of PLV populations possibly linked to non-photosynthetic populations.

A central finding of this study is the highly modular nature of PLV seasonality. Of the seasonal PLV populations identified at SPOT, WGCNA resolved a surprisingly large number of chronotypes, indicating that recurrent annual behavior is distributed across many distinct temporal niches rather than a few broad seasonal classes. This pattern is consistent with the growing realization that marine viral communities, including both bacteriophages and giant viruses, often exhibit repeatable interannual succession while retaining substantial fine-scale heterogeneity within the broader seasonal cycle. At SPOT, T4-like myoviruses were previously shown to display recurring spring-summer versus fall-winter patterns, alongside persistent moderately abundant taxa that remained detectable across much of the year (Dart et al., 2023). The SPOT bacteriophage community is itself known to exhibit strong seasonality, with approximately one-third of viral populations showing significant annual periodicity (Cram et al., 2016). More recently, monthly viral metagenomes from the Western English Channel identified multiple viral “chronotypes” with distinct annual trajectories and environmental associations (Bolaños et al., 2024), reinforcing the idea that temporal partitioning is a pervasive organizing principle of marine viral communities. Our PLV results extend this logic to a major but understudied group of eukaryotic DNA viruses.

The specific shapes of the dominant modules suggest that PLVs at SPOT are tracking several different recurring ecological windows. Some modules peaked in late autumn to early winter, others showed broader winter enrichment, and still others were strongest in late summer or early fall. This spread of seasonal peaks support a scenario in which distinct PLV populations are linked to different protist hosts, host physiological states, or infection modes that recur at predictable times of year. This temporal partitioning implies that PLV-host interactions are not synchronously driven by a single seasonal cue but are instead governed by a suite of partially independent environmental signals including different abiotic factors along with the timing of specific host lineage blooms, that together produce the mosaic of seasonal niches observed. Such a model is possible given the well-documented seasonality of microbial eukaryotes in marine systems. At SPOT, while chlorophyll and primary production show strong seasonal cycles, protistan communities are often less temporally stable than prokaryotic communities (Laperriere et al., 2025; Yeh & Fuhrman, 2022b). In that context, the many PLV chronotypes identified here likely reflect the highly dynamic protist communities in this system.

In bacteriophage systems, kill-the-winner dynamics predict that viruses should rise in response to expanding host populations and thereby contribute to cyclical suppression of dominant taxa. Similar ideas have increasingly been invoked for eukaryote-infecting viruses (Martínez et al., 2007; Pound et al., 2020), especially in systems where protist blooms or seasonal phytoplankton successions generate predictable host windows. However, PLVs are unusual because they seem to span a broad range of life strategies, with many PLV lineages known to integrate in host genomes, and some PLV lineages showing giant-virus-associated parasitism (C. Bellas et al., 2023; Stough et al., 2019). These factors are important to consider when interpreting the highly dynamic “chronotypes” observed here. Many PLV lineages may follow atypical life strategies compared with other DNA viruses of protists, including dynamics shaped by giant virus-induced activation that could generate rapid bursts in the abundance of specific lineages. In addition, environmental triggers may induce active virus production from integrated states in certain PLV lineages, further contributing to the temporal variability observed in this system. Thus, the seasonal modules described here may reflect not only the timing of host availability but also shifts in the relative prevalence of distinct life strategies, including productive infection, persistence, and possibly giant-virus-dependent replication in subsets of the PLV community.

The partial phylogenetic structure of seasonal modules is particularly informative in this regard. Several chronotypes were enriched for specific clades, especially PLV2, suggesting that some aspects of temporal behavior are conserved within lineages, possibly because related PLVs share host lineages, infection strategies, or genome-encoded functions that bias them toward similar temporal niches. However, this phylogenetic signal was incomplete as the same broad clade could appear in multiple temporal modules. This pattern demonstrates that phylogeny alone doesn’t dictate the seasonality of PLVs. In other words, evolutionary history constrains, but does not fully determine, ecological dynamics. A similar conclusion was recently reached for marine giant viruses and cyanophages from the same regional time series, where temporal succession was structured in part by phylogenetic proximity, but phenological divergence among close relatives still occurred, likely reflecting host shifts or ecological specialization (Dart et al., 2023; Laperriere et al., 2025). Our PLV results now suggest that this combination of phylogenetic constraint and ecological flexibility may be a broader property of eukaryote-infecting DNA viruses in marine systems. Finally, the roughly 50% of PLV populations that showed no significant seasonal signal (Cluster 1 in our analysis), may represent populations subject to stochastic boom-bust dynamics tied to transient physical or biological events or populations tracking hosts with highly variable bloom timing. Although viruses are generally very specific for their hosts, PLVs with persistent patterns may also represent populations with broad host ranges that can move among multiple temporally shifting hosts.

Co-occurrence analysis between PLV and eukaryotic abundance profiles provides a first-pass framework to infer potential host associations, although correlation-based approaches are inherently limited for this purpose. While correlations cannot establish direct infection, prior community-scale studies have shown that established host–virus relationships can often be recovered when sufficient temporal sampling is available. Leveraging the extensive 18S rRNA-based eukaryotic dataset from matched SPOT time points, we identified moderate to strong associations (Pearson’s r > 0.4) linking multiple PLV lineages to a diverse assemblage of eukaryotes. These include repeated associations of Trimcap/X PLVs with dinoflagellates, ochrophytes, and ciliophora, as well as links between PLV2 and radiolaria and between PLV1 and chlorophyta, suggesting that several PLV clades may interact with both phototrophic and heterotrophic protists. Notably, while most inferred associations involve protistan taxa, we also observe recurrent connections between specific PLV clades (particularly X group) and metazoans, including multiple independent links converging on the same metazoan node. Such associations are consistent with emerging evidence that polintons, which are evolutionarily related to PLVs, are widespread in metazoan genomes, including in marine animals such as corals. This raises the possibility that at least some PLVs may infect, or interact with, Metazoan hosts. More broadly, the frequent associations with ecologically central protist groups such as dinoflagellates, ochrophytes, and choanoflagellates point to a potentially important role for PLVs in structuring marine food webs and influencing biogeochemical processes through impacts on both primary producers and grazers. At the same time, these inferred links remain hypotheses that will need further validation. Nonetheless, the consistency of these associations across a multi-year, high-resolution time series lends confidence that they reflect biologically meaningful relationships and provides a focused set of testable host candidates of marine PLVs for future experimental and genomic validation.

### Functional repertoire of SPOT PLVs

Although most PLVs encode the expected structural and replication genes, many also contain genes with potential metabolic or host-modulatory roles, including ribonucleotide reductases, glycosyltransferases, lipase-like proteins, nucleotide-sugar transferases, glutaredoxin-related proteins, and other enzymes. Importantly, these genes are unevenly distributed across clades, and t-SNE clustering based on functional repertoires broadly recapitulates phylogenetic structure. That combination of lineage-specific gene content and ecological differentiation suggests that PLV diversification has involved the assembly of distinct functional repertoires that may shape infection phenotypes. In other viral systems, auxiliary metabolic genes are often interpreted as mechanisms for redirecting host metabolism toward conditions favorable for viral replication (Tian et al., 2024). However, caution is warranted here. The presence of a metabolic gene in a PLV genome does not by itself demonstrate that the gene is expressed, functional, or adaptive in the infection context. Instead, our data support the more conservative conclusion that some PLV lineages possess the genomic potential to modulate host carbohydrate, nucleotide, lipid, or redox processes, and that this potential is distributed non-randomly across the PLV phylogeny. Among these functions, glycosyltransferases and related carbohydrate-processing genes are interesting because they were especially common in Group X compared to other PLV groups. In large DNA viruses, glycosylation-related genes can influence host surface interactions, virion structure, or manipulation of cellular metabolism, although their precise roles might vary across viral lineages (Speciale et al., 2022). In contrast to carbohydrate-processing genes, ribonucleotide reductases were more common in PLV2 lineage. Ribonucleotide reductases are widely encoded in viruses that can enhance nucleotide supply during replication and are common in phages and giant viruses. The enrichment of RNRs in PLV2 therefore raises the possibility that some PLV lineages are better equipped for productive replication under conditions where nucleotide availability constrains infection.

## Conclusion

PLVs represent an emerging group of dsDNA viruses of protists and may contribute to marine food-web and biogeochemical dynamics more than is currently appreciated. Our results position PLVs as a major and ecologically differentiated component of the marine virosphere. At SPOT, PLVs are abundant, phylogenetically diverse, and structured into many recurring temporal modules. Their seasonality is partly shaped by phylogenetic history, but not reducible to it, indicating that ecological divergence occurs both within and among PLV clades. Their positive associations with productivity-linked environmental variables suggest that many are connected to the seasonal dynamics of microbial eukaryotes, while their functional repertoires point to lineage-specific capacities for host interaction and metabolic modulation.

More broadly, these findings reinforce the view that PLVs are not rare, enigmatic elements at the boundary of viruses and mobile elements, but active ecological players whose diverse life strategies may influence the population biology, evolution, and seasonal restructuring of marine protist communities.

## Methods

### Sample collection and filtration

Seawater for DNA samples was collected as part of the San Pedro Ocean Time-series. In this project we used monthly samples collected from May 2009 to September 2014. DNA samples from the viral fraction were obtained by filtering water onto a 0.22µm Sterivex cartridge followed by a 25 mm 0.02µm Antop filter cartridge.

### Sequencing

Viral fraction metagenomes were sequenced at the Department of Energy (DOE) Joint Genome Institute (JGI). Libraries were prepared from 1 ng of viral fraction DNA per sample according to manufacturer’s instructions (Swift 1S Plus or Nextera XT; Individual sample details are in the JGI Genome Portal under proposal 2799).

### Sequence processing/assembly

Raw Read data was retrieved from the JGI Genome Portal along with assemblies generated for these reads.

### Polinton-like virus identification and phylogenetic assessment

Proteins were predicted from megahit-assembled contigs using prodigal with the -p meta flag. PLVs were then identified using hmmsearch (Eddy, 2009) with an e value cut off of 1e-5 using hmm profiles for the known forms of the PLV major capsid protein (MCP) (C. M. Bellas & Sommaruga, 2021). Contigs containing a MCP hit over the e value threshold were clustered at 95% ANI similarity using cd-hit-est (Fu et al., 2012). The Longest Contigs from each bin was used as a population representative for phylogenetic analysis. The MCP proteins from these representatives were aligned with reference genomes using mafft (Katoh & Standley, 2013) with the –auto flag. A maximum likelihood tree was then assembled using IQtree (Nguyen et al., 2015) with 1000 Ultrafast Bootstraps, and visualized usi*ng iTOL (Letunic & Bork, 2024)*.

### Seasonal dynamics analysis

To assess the seasonal dynamics of the identified PLV viral species in this data set, raw reads obtained from the JGI Genome Portal were mapped to the identified PLVs using CoverM (Aroney et al., 2025). The resulting read mapping files were clustered based on seasonal variance using a Fourier Transformation from tslearn (Tavenard et al., 2020) library in python. With an optimized number of clusters determined based on silhouette score. These clusters were visualized using matplotlib (Hunter, 2007) to generate clustering graphs and a polar plot for each population. The statistical analysis of these clusters was performed using scipy.stats (Virtanen et al., 2020) to perform a Kruskal-Wallis, Friedman, and Ljung-Box test. Clusters that were identified to have seasonal trends if at least two tests showed a significant p value (<0.05).

### Functional clustering

Functional clustering was performed on a presence absence matrix of functional genes annotated using an in-house script with Pfam (Paysan-Lafosse et al., 2025) and VOG (Trgovec-Greif et al., 2024)databases. The presence absence matrix was used with a clade association dataset to generate an upset, bubble, and heatmap plot of the ten most abundant functional annotations. The upset, bubble, and heatmap plot were generated using R (R Core Team, 2021) in R studio (Posit team, 2025) with the upsetR (Gehlenborg, 2015) package and ggplot2 (Wickham, 2016).

### Environmental variable analysis

Relevant environmental variables for each sampling time point were obtained from the NCBI SRA. These variables were used in correlation analysis in python using scipy stats (Virtanen et al., 2020). Each variable was correlated with PLV abundance to determine their effect on the PLV populations over time was graphed using matplotlib (Hunter, 2007) displayed in figure 2. The significance of these correlations was assessed using Mantel test.

### Genome annotation

PLV genomes had their proteins predicted using prodigal (Hyatt et al., 2010) followed by annotation of functions using an in-house script and the Pfams (Paysan-Lafosse et al., 2025) and VOG (Trgovec-Greif et al., 2024) databases. A selected set of these annotated viruses were then visualized using GBdraw (Satoshikawato, n.d.).

### WGCNA/Chronotype analysis

The genome read count data generated in the initial seasonal dynamic analysis above was also used to generate a WGCNA network using the WGCNA package (Langfelder & Horvath, 2008) in R (R Core Team, 2021). The WGCNA network clustered genomes into chronotypes. Chronotypes were used to group genomes by seasonal trend as a method to assess the relationships between the viruses in question.

### Virus host association

The abundance profiles of the PLV groups identified in this study were analysed for co-occurrence against 18S abundance profiles from the cellular fraction. This association network was constructed using FlashWeave (Tackmann et al., 2019). The resultant connections were filtered and connections between PLVs and 18S profiles with positive weighting values greater than 0.4 were retained. The resultant network was visualized using Cytoscape.

## Data availability

The population representing contigs, raw phylogenetic tree, functional annotations and additional files are available at Figshare: https://figshare.com/s/31af6f9451480046926f

## Acknowledgements

M.M. and E.M. are funded by the Rosenstiel School of Marine, Atmospheric and Earth Science.

This work was also supported by the Simons Foundation (CBIOMES-549943 to J.A.F.) and the National Science Foundation (NSF-OCE-1737409 to J.A.F.). We thank the USC Wrigley Institute for Environmental Studies for their support of the SPOT sampling, Julio Cesar Ignacio Espinoza for his part in virus DNA extractions and sequencing library preparation, and the Department of Energy Joint Genome Institute for sequencing the DNA as part of their CSP program (proposal 2799 to N.A.A. and J.A.F.). C.B. was supported by a grant from the Austrian Science Fund (FWF P-34620)

