## Supplementary File for "Expansive Diversity and Temporal Dynamics of Emerging Polinton-like Viruses in a Marine Ecosystem"

Supplementary material for Mimick et al., 2026. This file contains legend for supplementary dataset 1, supplementary figures 1-5, and supplementary table 1.

**Supplementary Dataset 1 legend:** Correlation values (Pearson’s *r*) between the SPOT PLV and eukaryotic taxa represented in the association network (Figure 6). Only connections with *r>0.4* are shown.

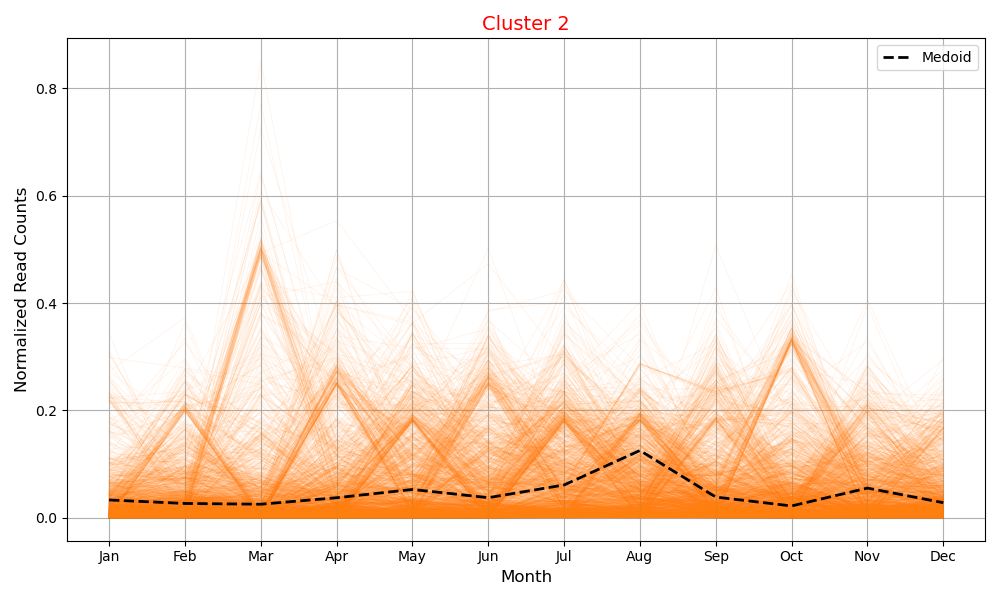

**Supplementary Figure 1:** (A) Relative abundance profile of the PLVs that are not seasonal in nature (cluster 2) (p>0.05). Orange lines show relative abundance of individual PLV populations over the months, and the black dashed line shows the median abundance of all the PLV populations represented in this plot. Also see Figure 3A for the PLV cluster that is seasonal.

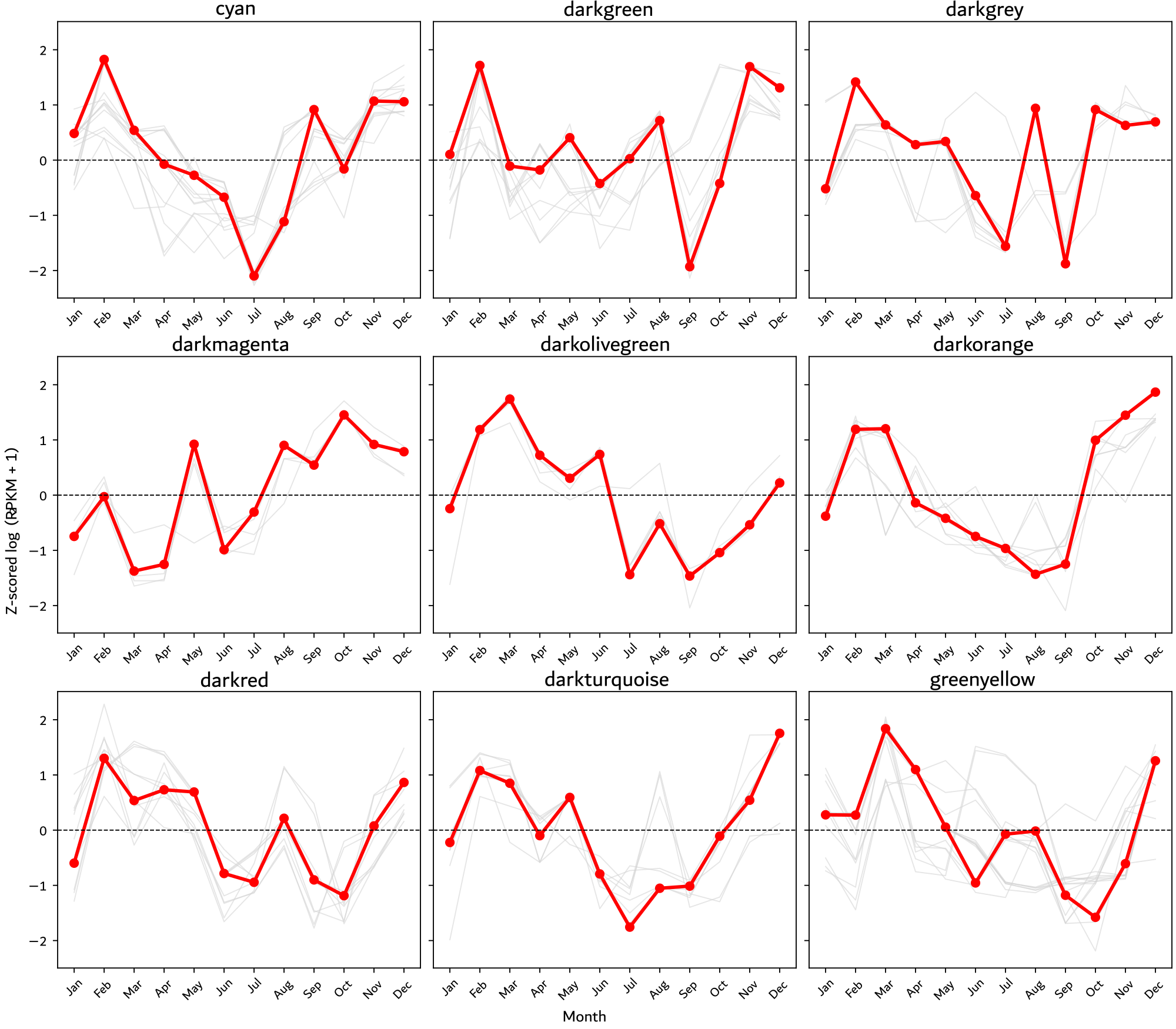

**Supplementary Figure 2:** Monthly abundance profiles of significantly seasonal PLV populations grouped into WGCNA modules. Nine modules are shown here - cyan, darkgreen, darkgrey, darkmagenta, darkolivegreen, darkorange, darkred, darkturquoise, and greenyellow. Gray lines represent individual PLV populations, while red lines show the average seasonal trajectory for each module, revealing distinct annual abundance patterns among PLV groups. For the trends of modules with the highest number of PLV populations, see Figure 3.

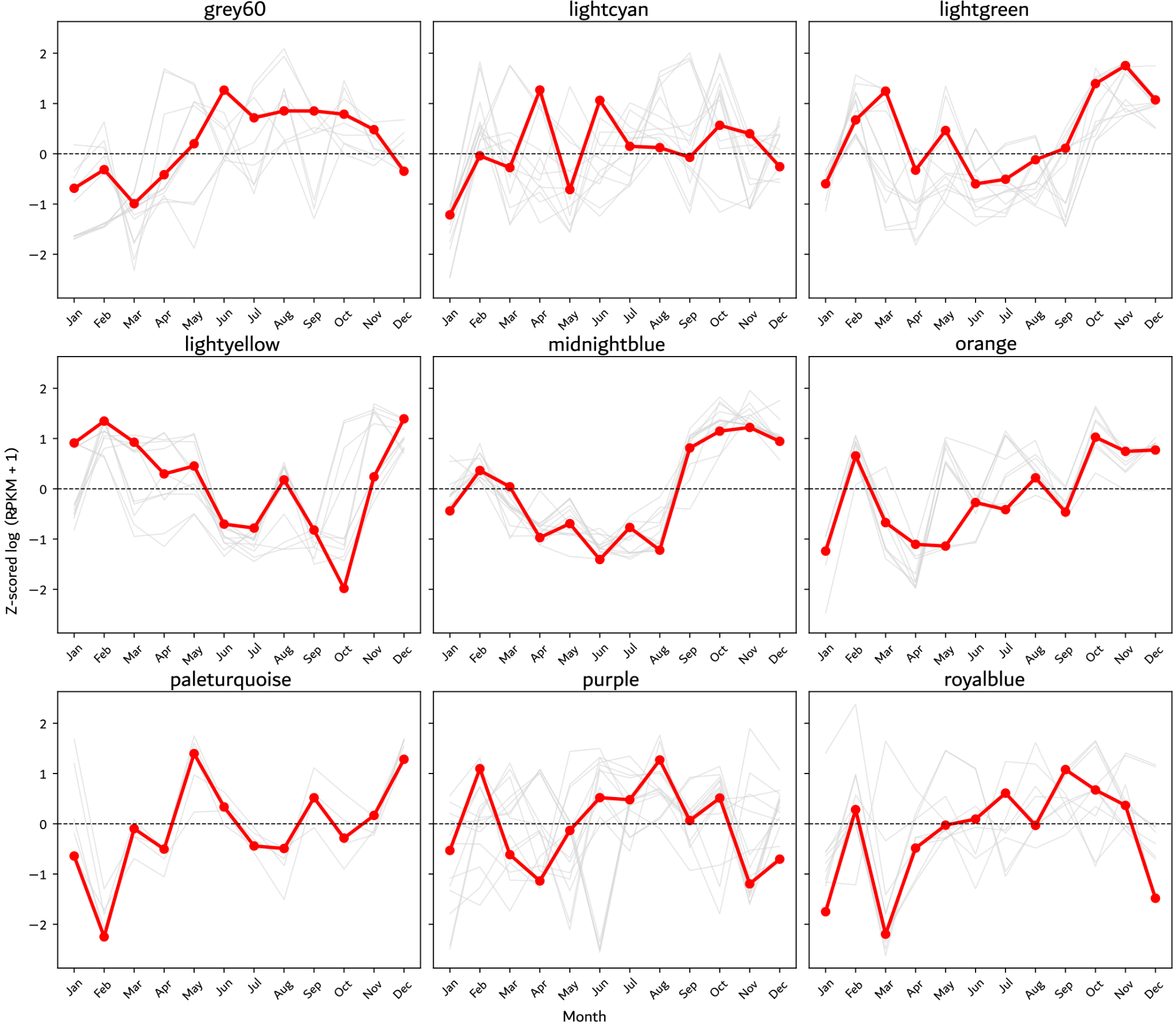

**Supplementary Figure 3:** Monthly abundance profiles of significantly seasonal PLV populations grouped into WGCNA modules. Nine modules are shown here: gray60, lightcyan, lightgreen, lightyellow, midnightblue, orange, paleturquoise, purple, and royalblue. Gray lines represent individual PLV populations, while red lines show the average seasonal trajectory for each module, revealing distinct annual abundance patterns among PLV groups. For the trends of modules with the highest number of PLV populations, see Figure 3.

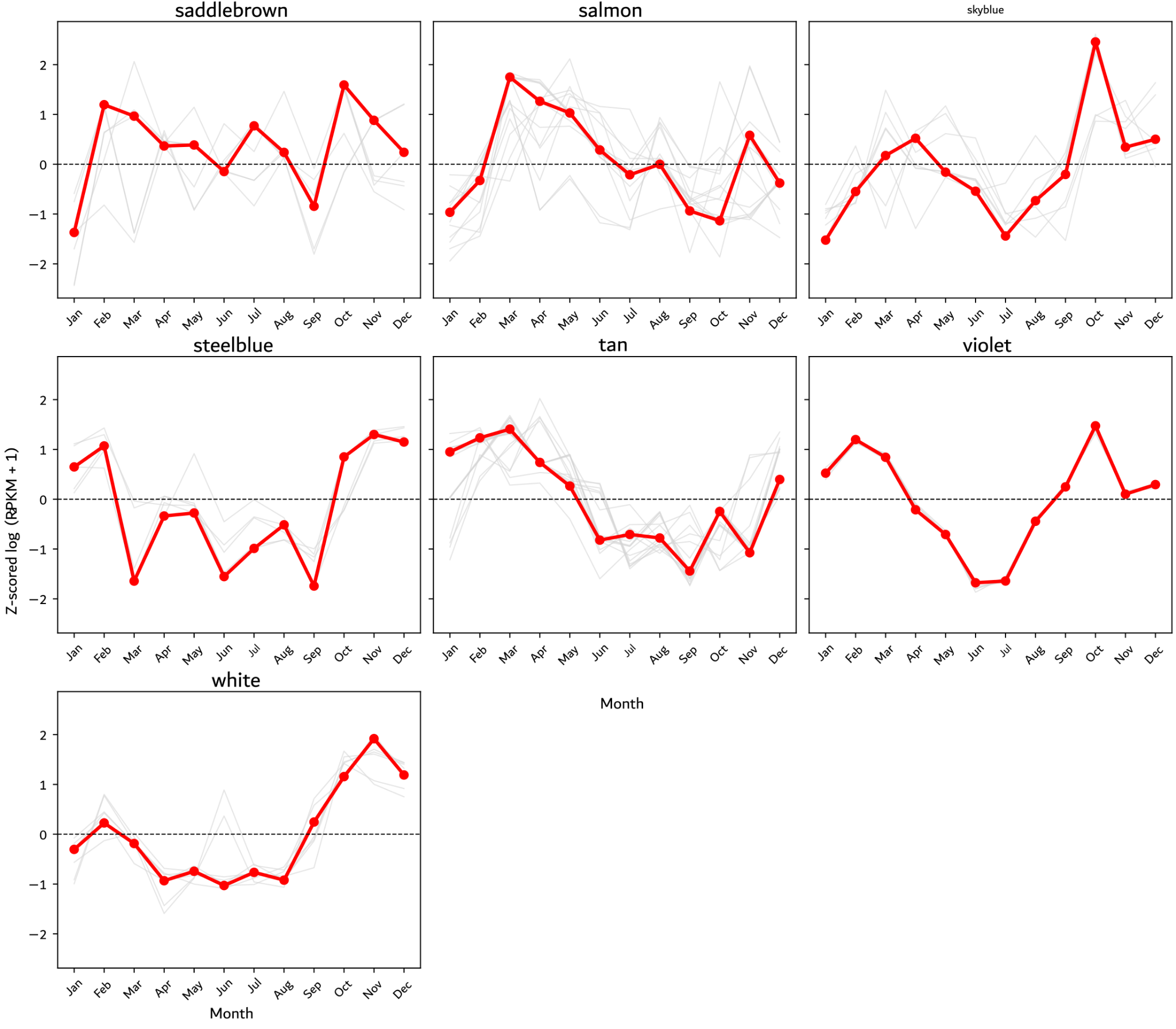

**Supplementary Figure 4:** Monthly abundance profiles of significantly seasonal PLV populations grouped into WGCNA modules. Seven modules are shown here - saddlebrown, salmon, skyblue, steelblue, tan, violet, and white. Gray lines represent individual PLV populations, while red lines show the average seasonal trajectory for each module, revealing distinct annual abundance patterns among PLV groups. For the trends of modules with the highest number of PLV populations, see Figure 3.

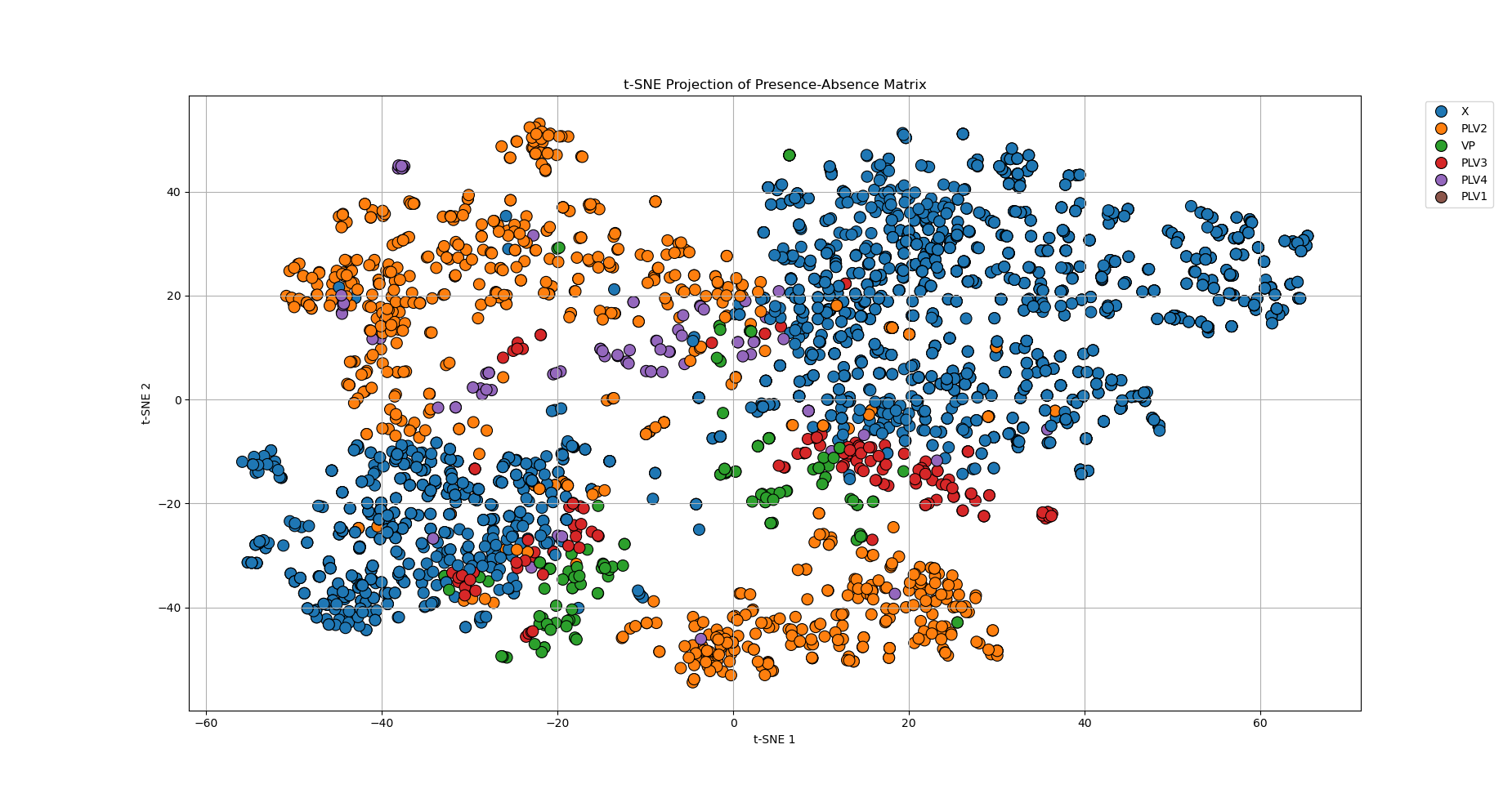

**Supplementary Figure 5:** t-SNE grouping of the functional patterns in the PLV clades at SPOT site.

**Supplementary Table 1**: Number of genomes per WGCNA module

| **module** | **count** | **module** | **count** |
| --- | --- | --- | --- |
| grey | 69 | lightgreen | 12 |
| turquoise | 41 | lightyellow | 11 |
| blue | 30 | royalblue | 10 |
| brown | 23 | darkred | 10 |
| yellow | 22 | darkgreen | 10 |
| green | 20 | darkturquoise | 9 |
| red | 19 | darkgrey | 9 |
| black | 18 | orange | 9 |
| pink | 17 | darkorange | 8 |
| magenta | 16 | skyblue | 7 |
| purple | 15 | white | 7 |
| greenyellow | 15 | saddlebrown | 6 |
| tan | 15 | darkolivegreen | 5 |
| cyan | 14 | paleturquoise | 5 |
| salmon | 14 | violet | 5 |
| midnightblue | 13 | darkmagenta | 5 |
| lightcyan | 13 | steelblue | 5 |
| grey60 | 12 |  |  |
